# Genomic evolutionary analysis in R with geaR

**DOI:** 10.1101/2020.08.06.240754

**Authors:** Christopher M. Ward, Alastair J. Ludington, James Breen, Simon W. Baxter

**Affiliations:** School of Biological Sciences, University of Adelaide, Australia; Bioinformatics Platform, South Australian Health & Medical Research Institute (SAHMRI), Australia; Robinson Research Institute, University of Adelaide, Australia; Faculty of Health & Medical Sciences, University of Adelaide, Australia; Bio21 Institute, School of BioSciences, University of Melbourne, Australia

**Keywords:** Evolution, Population Genomics, R Package, Admixture

## Abstract

The analysis and interpretation of datasets generated through sequencing large numbers of individual genomes is becoming commonplace in population and evolutionary genetic studies. Here we introduce geaR, a modular R package for evolutionary analysis of genome-wide genotype data. The package leverages the Genomic Data Structure (GDS) format, which enables memory and time efficient querying of genotype datasets compared to standard VCF genotype files. geaR utilizes GRange object classes to partition an analysis based on features from GFF annotation files, select codons based on position or degeneracy, and construct both positional and coordinate genomic windows. Tests of genetic diversity (eg. *d*_*XY*_, *π, F*_*ST*_) and admixture 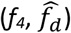 along with tree building and sequence output, can be carried out on partitions using a single function regardless of sample ploidy or number of observed alleles. The package and associated documentation are available on GitHub at https://github.com/CMWbio/geaR.

## Introduction

Improvements in genome sequencing technologies has led to increased production of data at lower relative cost per base (Schwarze et al. 2020). Genome-wide sequencing datasets with hundreds of samples can be produced for population genomic analysis, allowing researchers to investigate population and evolutionary history at an unprecedented scale. However, due to file size and data complexity downstream problems during data storage and analysis can arise. The most common format for handling genome-wide SNP data is the Variant Call Format (VCF), which has historically had a large memory overhead when being read into an R environment. To resolve this, the Genomic Data Structure (GDS) format has allowed all genotype and metadata to be compressed into a queriable, on-disk file that substantially reduce memory requirements and decrease analysis time (Zheng et al. 2017). The GDS format provides an efficient format for filtering SNP data in order to perform Principal Component Analysis, estimate genetic relatedness and tests for genetic association (Zheng et al. 2012).

GDS files use GRange objects from the GenomicRanges package (Lawrence et al. 2013) to define loci to query from file and import into R. In their most basic form, GRange class objects define genomic loci based on reference position. Although widely used throughout Bioconductor, GRange objects, to our knowledge, have not been utilized in the same manner to define loci for evolutionary analyses.

Few R packages attempt to carry out genome-wide investigation of genotype data. Most packages focus on the analysis of single or multi-locus data, with the notable exception of PopGenome (Pfeifer et al. 2014). However, one limitation of PopGenome is customizability of how the target genome is partitioned, and which sites are selected for analysis. Most tools, including PopGenome, allow datasets to be partitioned into sliding or tiled windows based on reference or SNP position. PopGenome also provides methods to split data into GFF attributes, however selection of bespoke partitions not possible. This makes calculating population metrics on specific codon positions (eg. four-fold or zero-fold degenerate sites) or analysing many non-contiguous loci difficult and time consuming.

To overcome these issues, here we present the R package geaR, which leverages the GDS format, to efficiently construct GRanges containing genome-wide or local loci of interest and to carry out common tasks for evolutionary analysis on genome-wide genotype data using a single function. Furthermore, we provide methods to partition the genome based on annotation, codon position or degeneracy through utilizing data in GFF files, or using reference genome coordinates or genotype position.

## Features

### Input data

Genotype input files are required to be in GDS format, enabling high compressibility compared to gzipped VCFs (>5X smaller on disk), efficient querying and the capability to work on large datasets with a reduced memory footprint (Zheng et al. 2017). Conversion of sample genotypes in the VCF format to a Genomic Data Structure (GDS) format can be performed using the SeqArray package (Zheng et al. 2017) before analysis with geaR. Genotypes called at any level of ploidy can be utilized in geaR, which includes whole genome sequence data generated from pools of two or more individuals.

### Partitioning the genome using GRanges

The geaR package utilizes GRange objects to define partitions for the analysis, for example, segmenting a genome into 10-kb windows. This allows users to define their own GRanges for the analysis or build them with provided functions. Currently, users are able to generate both coordinate (based on reference coordinate) and positional (based on genotype number) windows using *makeWindows()* or *makeSnpWindows()* functions. Sequence features, such as protein coding regions, can be extracted from a GFF with *getFeatures()*.

Many evolutionary analyses seek to calculate population metrics over different codon positions. To make this as simple as possible, geaR provides methods to index a reference genome according to codon position with *buildCodonDB()*, which can either be stored in memory as a GRangesList object or an SQLite database (DB) on disk to limit static memory usage. Users may then filter codons based on degeneracy (0-fold or 2-fold) and position using the function *filterCodonDB()*. A codon DB can also be passed to the function *validate4fold()* to select 4-fold degenerate sites across the genome that are empirically supported in the GDS file. This is done by querying the GDS to i) remove codons with missing data, ii) select 4-fold degenerate codons, iii) remove all those where codon positions one or two have variation and iv) select third positions.

GRange objects generated using geaR can then be combined using *mergeLoci()* to further customize partitions. For example, genome-wide tiled windows can be combined with four-fold degenerate sites to output either genomic windows that contain only 4-fold sites or all sites excluding 4-fold degenerate sites.

### Setting up an analysis: cogs and gears

geaR operates through two S4 classes, the ‘cog’ and ‘gear’ (Figure 1). Cogs, built using *makeCog()*, specify multiple analyses to carry out (see Table 1) setting parameters specific to each analysis. A single gear object can then be constructed, using *makeGear()*, which contains all of the specified cogs for analysis, along with the genomic loci and population metadata (Figure 2A). The *analyzeGear()* function then performs all analyses on the same set of genomic loci and samples, greatly reducing run time compared to sequential execution.

**Table 1:** Analysis types and functionality available to apply at each locus.

| Cog type | Functionality |
| --- | --- |
| Genetic diversity | Nucleotide diversity ($\pi$), genetic distance ($d_{xy}$), maximum distance ($d_{max}$), minimum distance ($d_{min}$), ancestral distance ($d_a$), $\gamma_{ST}$, relative node distance (RND), minimum relative node distance (RND<sub>min</sub>), and G<sub>min</sub> |
| Admixture | $f_4$ and $\hat{f}_d$ |
| Output loci | fasta format output to file |
| Output trees | newick format distance trees and metadata output to file |

**Figure 1:**
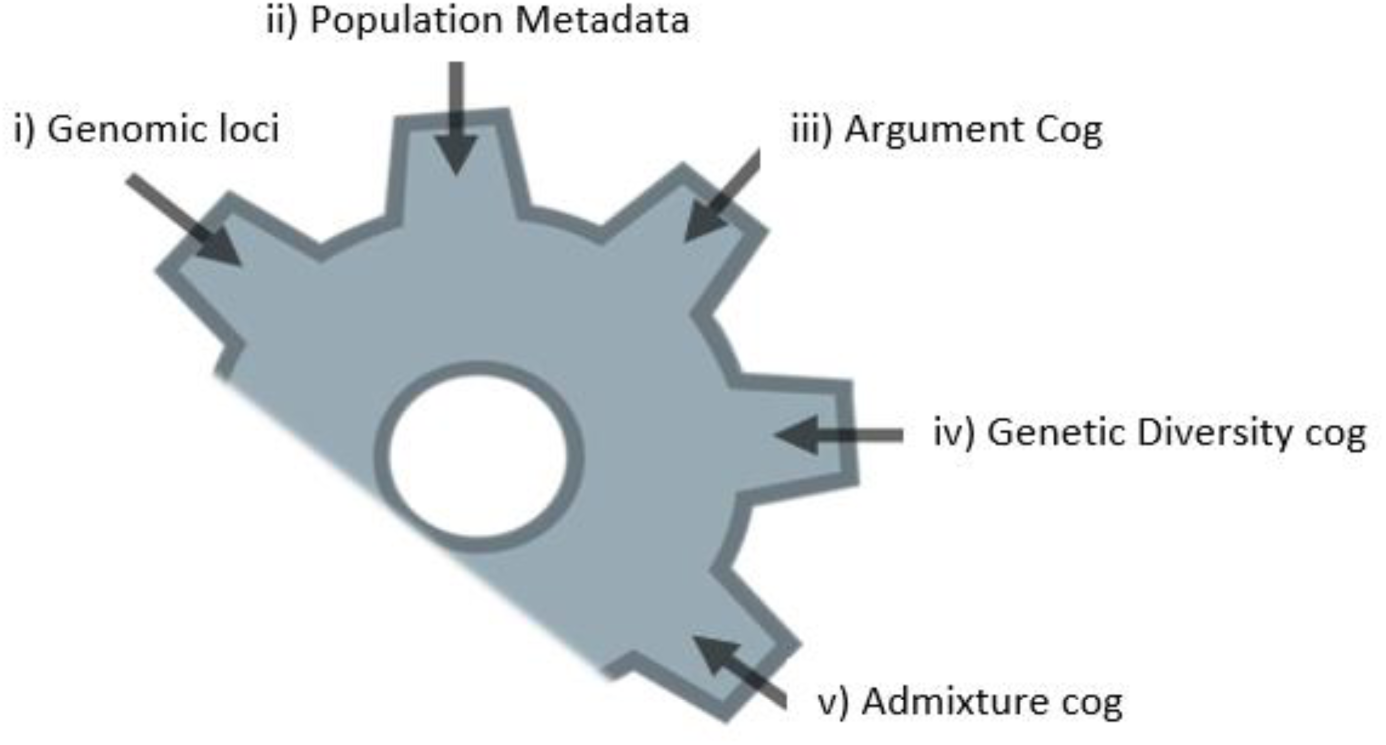
Structure of the of the gear S4 object: i) Genomic loci (GRanges) to carry out analyses across, ii) Population metadata encoding sample names to the population/species they belong to, iii) A cog containing general arguments for all analysis, iv) a cog specifying that the Genetic Diversity module should be carried out and v) a cog specifying that the Admixture cog should be carried out on the dataset.

**Figure 2:**
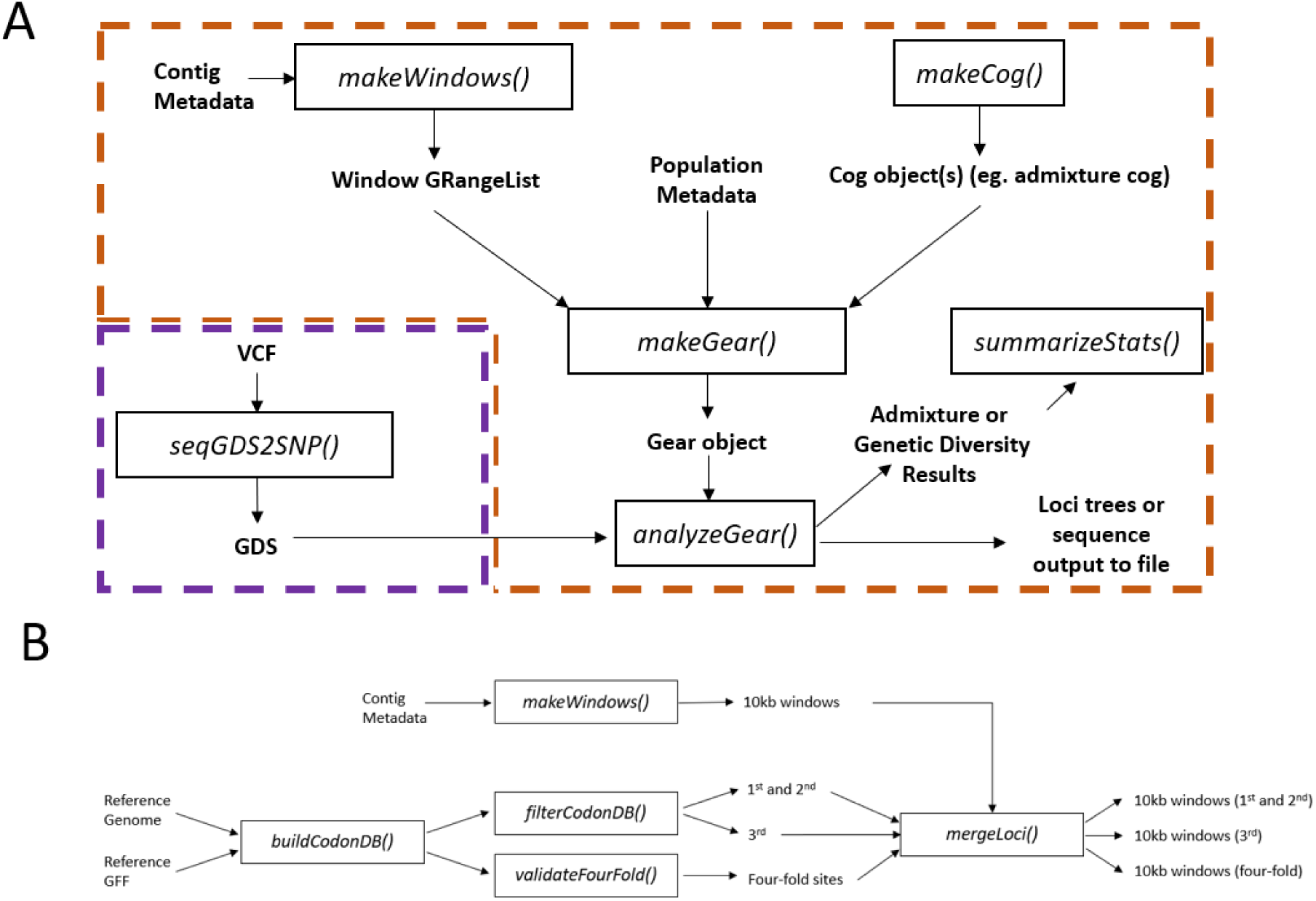
A) Basic analysis workflow in geaR to carry out analysis on windowed genomic loci. Functions specific to geaR are within the orange boundary and external functions in purple. After converting the VCF to GDS format using SeqArray, contig metadata (contig length) is used to construct windows across the genome. A dataframe containing population metadata defining population to sample grouping is then constructed. This is used, along with windows for the analysis and cogs, to construct the gear class object. The analysis is then carried out on the gear object and outputs depend on which cogs were specified. B) Workflow used to generate partition schemes for examples in Figure 3. First 10kb windows were generated from contig metadata. This was followed by generation of a codon database that indexes codon position in a reference genome. The codon database was then passed to the function *filterCodonDB* to output separate loci-sets for four codon partition schemes: 1^st^+2^nd^; 3^rd^; 0-fold and 2-fold. *<u>validateFourFold()</u>* was also used to select 4-fold degenerate sites that are supported by genotypes in the GDS file. Each of these codon loci-sets can then be passed to the function *mergeLoci()*, along with the 10kb windows, to combine loci into 10kb windows that contain only the selected codon types.

### Analysis types

Four different cogs can be generated to carry out an analysis: i) genetic diversity, ii) admixture, iii) outputLoci and iv) outputTrees. Genetic diversity allows the calculation of a range of population metrics (Table 1), most of which rely on genetic distance which is calculated based on the hamming distance between haplotypes at all sites within the locus. The admixture cog utilizes outgroup polarized allele frequency at all biallelic sites within the locus to calculate *f*_*4*_ (Patterson et al. 2012) and 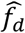 (Martin et al. 2015) statistics. The package also enables users to output data in fasta format for each individual (or sample pool) using outputLoci or as distance trees using outputTrees. Haplotypes are used in diversity calculations and are output to file according to the phase within the supplied GDS file, not calculated by geaR.

Outputs of both genetic diversity and admixture cogs can be summarized using *summarizeStats()* which calculates a mean and median values for each statistic across all loci using a block jack-knife approach.

### Parallelization

All functions allow operations to be run in parallel by leveraging methods in the furrr (https://github.com/DavisVaughan/furrr) and parallel R packages.

## Carrying out an analysis using gear

geaR has successfully been used to calculate genome wide diversity metrics between populations containing 532 moth genomes (You et al. 2020) and to identify introgressed regions between two *Bactrocera* fly species (Ward et al. 2020). Below we outline two example analyses using geaR. Code for each of examples and other common workflows can be found on the wiki (https://github.com/CMWbio/geaR/wiki).

In our first example we use a subset of the data from Ward and Baxter (2018) containing three populations of diamondback moth collected from Australian Capital Territory, Australia; South Australia, Australia and Hawaii, USA. Second. Following the general workflow shown in Figure 2A, we converted the called genotypes to the GDS format using SeqArray, constructed partitions, built cogs, combined those cogs into a gear and then carried out the analysis. We constructed our partitions for the analysis by generating a GRange object containing only scaffold_4. The analysis will use this GRange object to construct six different partition schemes based on 10kb tiled windows (workflow shown in Figure 2B): i) all sites, ii) only 1^st^+2^nd^ codon positions, iii) windows only 3^rd^ codon positions, iv) only 0-fold degenerate sites, v) only 2-fold degenerate sites and vi) only 4-fold degenerate sites. Partition i) was then used to calculate pairwise genetic distance (d_XY_) between each population across the scaffold (Figure 3A) and partitions ii-vi) were used to calculate within population nucleotide diversity across the whole genome (Figure 3B).

**Figure 3:**
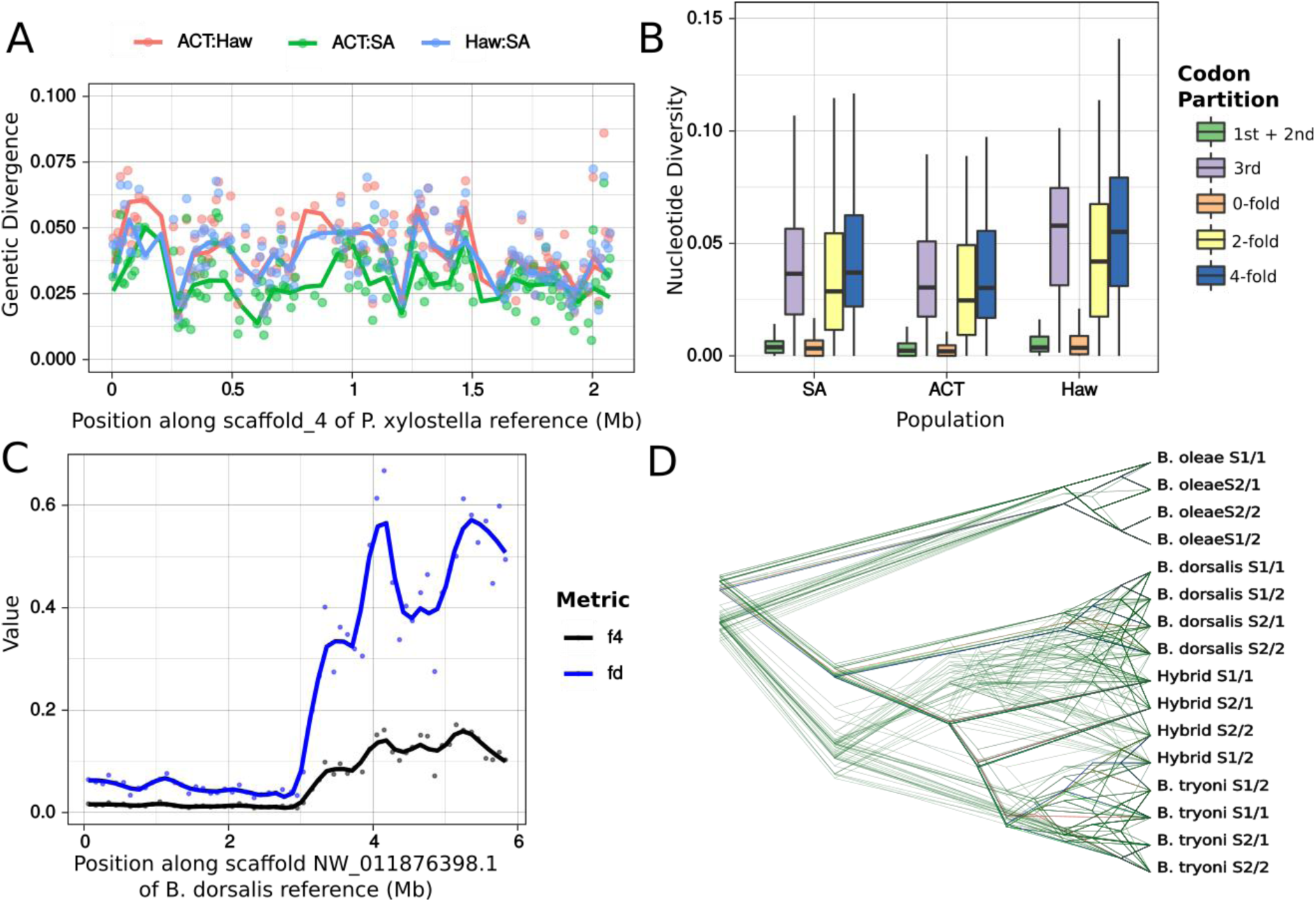
Example workflows to carry out with geaR: Panels A and B use *P. xylostella* data from Ward and Baxter (2018), panels C and D use *Bactrocera* from Ward *et al* (2020). A) Absolute genetic distance (*d*_*XY*_) was calculated between pairwise comparisons of three populations for 10kb tiled windows across scaffold_4 of the diamondback moth reference genome. C) The five loci-sets constructed using the workflow in Figure 2B were used to calculate nucleotide diversity (π) at 1^st^+2^nd^, 3^rd^, 0-fold, 2-fold and 4-fold codon sites across scaffold_4 of the diamondback moth reference genome C) Admixture metrics *f*_*4*_ and 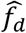 calculated on 100kb windows across scaffold NW_011876398.1 of the *B. dorsalis* reference genome. D) Distance trees for each 100kb window across NW_011876398.1 output using the *outputTrees* cog showing a mixture of discordant and concordant topologies. Plots A), B) and C were generated using ggplot2 (Wickham 2009) and D) using densitree (Bouckaert 2010).

For a second example we will identify one of the introgressed regions from Ward et al. (2020). This will use data from a single scaffold (NW_011876398.1) of the *B. dorsalis* reference genome (GCF_000789215.1) for two samples of *B. tryoni, B. dorsalis, B. oleae* and a *B. dorsalis/B*.*tryoni* hybrid line. Using the same methodology as the first example, we constructed a 100kb tiled window partition scheme. However, for this analysis we used the admixture cog to calculate *f*_*4*_ and 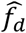 admixture metrics showing clear evidence for introgression at the 3’ end of the scaffold (Figure 3C). We also used the outputTrees cog to output distance trees for each of these windows to illustrate both the congruent and incongruent topologies resulting from partial admixture on NW_011876398.1 (Figure 3D).

## Conclusion

Genome-wide datasets with many individuals are becoming the norm in population genetic studies, increasing the need for tools to efficiently carry out analyses on genotype data. The functional programming capabilities of the R programming language provide an intuitive environment for users to carry out calculation and visualization of population and evolutionary genomics metrics. The methods provided in geaR allow users easily and effectively partition the genome for generic and bespoke analysis of genome-wide genotype data regardless of sample ploidy and number of observed alleles.

## Acknowledgements

We would like to thank Simon Martin for allowing us to use his python scripts as a reference for the first draft of this package. CMW is funded by the Commonwealth Hill Trust and The Grains Research Development Corporation (Grant 9175870).

## Data Availability

All data is available in the referenced publications.

## Notes

### Competing Interest Statement

The authors have declared no competing interest.

